# Increasing the shelf life of tomato fruit by editing the β-D-N-acetylhexosaminidase (β-hex) gene using CRISPR/Cas9 technology

**DOI:** 10.64898/2026.05.01.722371

**Authors:** Anvarjon A. Murodov, Mirzakamol S. Ayubov, Mukhammadjon Kh Mirzakhmedov, Nurdinjon S. Obidov, Bekhzod O. Mamajonov, Abdurakhmon N. Yusupov, Ziyodulloxon H. Bashirxonov, Lola K. Kamalova, Shukhrat O. Kushakov, Ilkhomjon E. Bozorov, Zabardast T Buriev, Ibrokhim Y. Abdurakhmonov

## Abstract

Obtaining tomato plants with firm and intact fruit is one of the main goals in tomato breeding programs. Achieving these goals through conventional breeding is time-consuming and can lead to the loss of unwanted traits. In other hand, consumers are concerned about the presence of transgenic elements in plants acquired through RNA interference. The use of CRISPR/Cas9 technology has made it possible to overcome the above-mentioned shortcomings. In this study, the β-D-N-acetylhexosaminidase (*β-hex*) gene, which is involved in tomato fruit ripening, was knocked out using CRISPR/Cas9. In the resulting mutant plant genome, an indel mutation was found in exons 1 and 2 of the *β-hex* gene. Plants with a mutation in their genome were observed to have increased fruit firmness and shelf life compared to control plants without affecting fruit quality.

## Introduction

Tomato (*Solanum lycopersicum* L) is one of the most widely grown vegetables. In 2024, 192.3 million tons of tomatoes were produced worldwide (FAO, 2024). Tomato fruit is rich in vitamins A and C, minerals, and antioxidants, which are important for maintaining human health [1]. Tomato is also important as a model plant for studying the processes that occur during the ripening of climatic fruits. It is also studied by scientists as a model plant, due to the complete genome sequence of several of its variants and the ease of genetic transformation [2].

50 % of the harvested tomatoes in the developing countries lose their quality before they reach consumers [3]. The main reason for this is that fruits that have lost their resistance to post-harvest damage lose their skin integrity as a result of external mechanical influences, and as a result, they are more susceptible to bacterial and fungal diseases [4]. To overcome this problem, scientists have used various physical, chemical, and genetic engineering methods to increase the shelf life of tomato fruit [5]. However, the application of certain physical and chemical preservation methods has been reported to induce significant alterations in the flavor profile and nutritional composition of the fruits [6,7]. Genetic engineering methods are an alternative solution to solve such a problem. Fruit shelf life was significantly increased when the activity of genes involved in fruit ripening, such as 2C protein phosphatases (*SlPP2C*) [8], aminocyclopropane-1-carboxylate (*ACC*) [9], and MADS-box protein (*SlCMB1*) [10] was suppressed by RNAi. However, public opinion remains concerned about plants modified through RNAi, especially those used for consumption [11]. One of the main reasons for this is that stable expression RNAi plants are GM [12]. In such circumstances, the use of CRISPR/Cas9 technology, a modern method of genome editing, has made it possible to reduce this concern by obtaining transgene-free organisms [13]. CRISPR/Cas9 also stands out from other genetic engineering methods in its ease, convenience, and low cost [14].

CRISPR/Cas9 has been widely used to improve fruit storage. In particular, variants with extended shelf life were obtained by knocking out the genes *CmACO1* in melon [15], *FIS1* and PL in tomato [16], *PG2a* [17], and *FaPG1* in strawberry [18] using CSIPR/Cas9.

More than 22 enzymes have been identified to work together during tomato fruit ripening [19]. Among these, direct softening of the fruit skin is observed as a result of the activity of enzymes that break down cell wall components [20]. The cell wall is mainly composed of cellulose, hemicellulose, pectin, and proteins. When the activity of the enzymes Polygalacturonase, β-galactosidase, and β-glucanase, which break down cell wall components, was inhibited through genetic engineering, only a slight increase in fruit shelf life was achieved [21–23]. The cell wall also contains N-glycoproteins, and the breakdown products of these substances, free N-glucans, are precursors for the process of glycosylation or glycoprotein proteolysis [24]. These free N-glucans are present throughout the entire developing pericarp of tomato fruit and increase during ripening [25].

β-D-N-acetylhexosaminidase (β-Hex; EC 3.2.1.52) is also one of the N-glucan hydrolyzing enzymes, and its function is to hydrolyze the last N-acetyl-D-hexosamine residues that are part of many N-glucans [26]. This enzyme is ripening specific, and its biosynthesis has been found to increase during fruit ripening [24]. Expression of beta-hex gene is regulated by RIPENING INHIBITOR (RIN), ABSCISIC ACID STRESS RIPENING 1 (*SLASR1*), and ethylene [27]. Post-harvest treatment of strawberry fruits with alginate oligosaccharides (AOS) resulted in a decrease in β-Hex enzyme activity, resulting in delayed fruit softening in the fruits [28]. It has been found that this enzyme increases in quantity during the ripening period of tomato fruit [25]. It has been observed that the shelf life of fruits is extended when the expression of the *β*-*hex* gene encoding this enzyme is reduced using RNAi technology [24]. Using site-directed mutagenesis, it was demonstrated that mutations within the cis-acting elements of the tomato *β*-*hex* promoter significantly downregulated its transcriptional activity, thereby delaying the fruit softening process and extending shelf life. Our earlier study [5] covered more details on various methods utilized by numerous laboratories to extend the shelf life of tomato fruits. In this research, we knocked out the *β*-*hex* gene using CRISPR/Cas9 technology and obtained transgenic free plants with extended shelf life.

## Materials and Methods

### Plant Materials and Growth Condition

The seeds of *S. lycopersicum* L. (“cherry” cultivar) were obtained from the Research Institute of Plant Genetic Resources, Uzbekistan. The surface of the seeds was sterilized by first soaking in 70% ethanol for 5 minutes, then in 3% sodium hypochlorite for 8 minutes. The seeds were then rinsed in sterile distilled water 3 times for 10 minutes each and stored on sterile filter paper until excess moisture was removed. The seeds were then grown in a growth chamber with a 16/8 h light-dark photoperiod and 70% humidity for 8-10 days on MS (Murashige and Skoog) medium containing 3% sucrose and 0.3% Gelzan (PhytoTech Labs, Inc. 14610 W. 106th St, Lenexa KS 66215). Fully formed 8-10-day-old cotyledons were used for transformation.

### sgRNA Design and CRISPR/Cas9 Construct Generation

sgRNAs of 19 bp in length were designed to target exons 1 and 2 of the *β*-*hex* gene using the online tool CRISPR-P 2.0 (http://crispr.hzau.edu.cn/cgi-bin/CRISPR2/CRISPR) [29]. sgRNA_1: GTCGCCGCAGATATGTAGG for exon 1 and sgRNA_2: GCCGCTATGATCAGCCACC for exon 2 were selected (S1 Table). Their minimum off-target mismatch is 3 bp and 4 bp and GC content is 55 and 60%, respectively (S2 Table). The secondary structure of gRNAs was determined using the web program RNAfold (“RNAfold Web Server,” n.d.). The construct for the CRISPR/Cas9 tomato transformation was assembled using the CasCade vector system [30].

### Agrobacterium-mediated transformation

The assembled vector construct was transformed into tomato plants using 8-10 day old cotyledons and *Agrobacterium tumefaciens* strain LB4404. The tops and bottoms of the cotyledons were cut off and infected with Agrobacterium (OD600=0.6) containing our vector construct. First, 2 days in the dark at 19 C in co-cultivation medium, then 2 weeks in 2Z selection medium1 at 24 °C ± 2 °C, with a 16-h photoperiod. Next, 1Z Selection medium 1 was maintained under the same nutrient conditions as above. The shoots (2 cm) that reached the top of the Magenta box were cut and maintained in selective rooting medium until their roots were sufficiently formed. Later it was adapted to the soil. We used hygromycin (6 mg/l) as a selective antibiotic. Plant transformation was conducted following the *Agrobacterium tumefaciens*-mediated method as previously described by Van Eck et al. (2019). [31].

### Molecular analysis of CRISPR-edited plants

To confirm the presence of the CRISPR/Cas9 construct and identify genomic mutations, genomic DNA was extracted from the young leaves of 8-week-old transplants using the Thermo Scientific™ GeneJET Plant Genomic DNA Purification Kit (K0791). The initial screening of T_0 generation plants was performed via PCR using Cas9-specific primers (S1 Table) to verify the integration of the genetic element. DNA from wild-type (WT) plants served as a negative control. The PCR thermal cycling conditions consisted of an initial denaturation at 95°C for 5 min, followed by 35 cycles of 95°C for 30 sec, 59°C for 30 sec, and 72°C for 40 sec, with a final extension at 72°C for 5 min. For mutation characterization in T_0 plants, PCR products amplified with gene-specific primers were first cloned into the TOPO™ TA Cloning™ Kit (Thermo Fisher Scientific, USA) due to the potential chimeric or heterozygous nature of the primary transformants. Subsequently, positive clones were sequenced using the SeqStudio Genetic Analyzer. In contrast, for the T_2 and T_3 generations, where mutations reached homozygosity, genotypes were determined through direct Sanger sequencing of the PCR products without intermediate cloning.

### Off-target analysis

“Potential off-target sites were identified using the CRISPR-P 2.0 online tool. Among these, site with high off-target score that was located within intron regions were selected for experimental validation. These loci were amplified via PCR and verified through Sanger sequencing to ensure the absence of unintended mutations (S2 Table).

### Protein prediction

The Expasy Translate program (https://web.expasy.org/translate) was used to predict the protein sequences of the mutants and the WT sequence.

### Fruit phenotyping

Phenotypic parameters such as fruit weight, fruit shape index (FSI), pH, and weight loss were evaluated in mutant and wild type plants. First, all fruits were sterilized in distilled water and ethanol. 10 fruits at the red ripe stage were used to determine fruit weight and fruit shape index (FSI).

To measure fruit shape, the ratio of the polar and equatorial diameters of the fruits was calculated. The pH was measured using a FiveEasy Plus Benchtop FP20 pH/mV Standard Kit (METTLER TOLEDO, Malaysia) in 3 biologicals and 3 technical replicates by diluting 250 mg of pericarp in 15 ml of distilled water (pH 7.0).

Fruit weight loss was also measured every 3 days for 8 fruits at the ripe red stage up to day 18. Weight loss was calculated using the following formula:

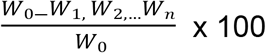

where W_0_ is the initial picked fruit weight, W_1_ is the fruit weight 3 days after harvesting, W_2_ is the fruit weight 6 days after harvesting, and so on up to n=6.

Also, the color change during the ripening process of the fruits was monitored every 5 days for a total of 15 days in the Breaker (Br) stage fruits.

### Data analysis

Statistical analyses were performed using GraphPad Prism 8.0 (GraphPad Software, San Diego, CA, USA). Data are presented as mean ± standard deviation (SD). The significance of differences between the wild-type (WT) and mutant lines was determined using Student’s t-test, with p-values < 0.05 considered statistically significant (*p < 0.05, **p < 0.01, ***p < 0.001; ns, non-significant).

For bioinformatic analysis, DNA sequencing chromatograms were visualized and aligned using SnapGene software (GSL Biotech; available at snapgene.com) and VectorBee (VectorBee.io). These tools were utilized to verify the presence of indel mutations, analyze frame-shift events, and ensure the integrity of the CRISPR/Cas9 vector constructs. Sequence translations and protein secondary structure predictions were further processed using the ExPASy Translate tool.

## Result

### Two mutation events in *β-hex* gene were generated by CRISPR/ Cas9 in “Cherry”

A total of 9 putative plantlets were recovered following Agrobacterium-mediated transformation. Molecular screening via PCR revealed that 3 plants carried the target construct. However, significant phenotypic abnormalities and premature senescence were observed in two transgenic lines, preventing them from reaching the reproductive stage. Similar phenomena regarding the low survival rate of primary plantlets (T0) have been widely reported in plant biotechnology, often attributed to tissue culture-induced stress or insertional mutations. As a result, only one vigorous T0 line was advanced to the T1 generation for segregation analysis (Fig 1C). Analysis of the plant harboring the vector construct using Sanger sequencing revealed a 1 bp insertion mutation in exon 1 and a 5 bp deletion mutation in exon 2. (Fig 1A).

**Fig 1.**
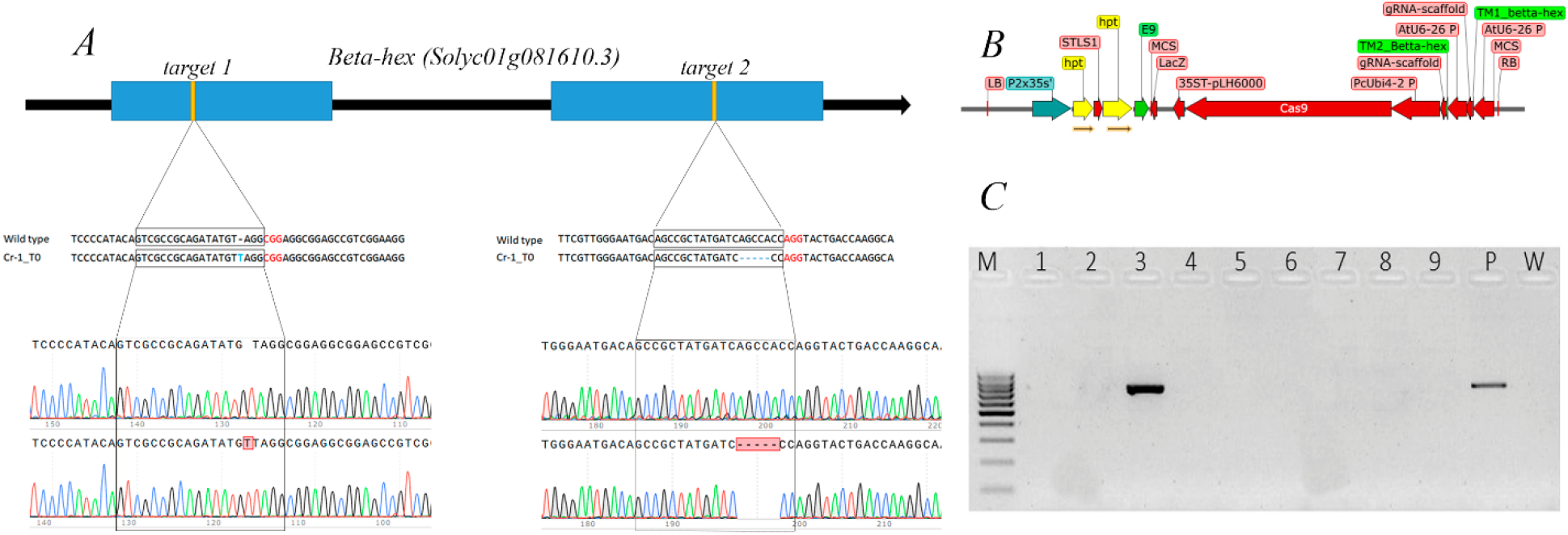
Vector construct structure and mutant variants in the resulting transgenic plants. **A** Location of sgRNAs in the *β-hex* gene, types of mutations produced, and Sanger sequencing chromatogram (PAM is shown in red and gRNAs are shown in boxes). **B** vector construction structure. **C** Presence of Cas9 construct in β-hex^in/del^ plant genome as seen on agarose gel (M-DNA ladder (100bp-1000bp С интол), 1-9 T0 plants, P positive control (plasmids), w –wild type).

The changes in the β-D-N-acetylhexosaminidase (β-Hex) protein resulting from these mutations were analyzed using the Expasy program. According to the results, it was predicted that a premature stop codon may occur in the sgRNA-located part of exon 1 and cause protein coding disruption. In wild type (WT) plants, the stop codon comes after the Serine amino acid at position 575, while in mutant plants, a premature stop codon is formed after the Alanine amino acid at position 127 at the sgRNA mutation site.

### Off-target analyses

Cas9 free plants were identified among the T_2 generation plants and off-target analyses were performed on these transgenic free plants. For 1-gRNA, 6 off-target cites with the highest off-target scores were identified, 2 of which corresponded to intron parts of the genome with 4 mismatches, and these 2 off-target parts were analyzed using Sanger sequencing. For 2-gRNA, sequencing was not performed because all of the high off-target scores corresponded to intergenic parts of the genome. According to the sequencing results, no mutations were detected in the off-target parts (S3 Fig).

### Protein prediction analysis

The effect of the mutations on protein coding was analyzed using the Expasy program. It was found that the insertion in exon 1 and the deletion in exon 2 resulted in the formation of premature stop codons. (S1 Fig).

### β-hex^in/del^ showed no difference in morphologic traits except late ripening and weight loss

A progressive linear increase in cumulative weight loss was observed across all samples over the storage period. This is a characteristic physiological process primarily driven by transpiration and respiratory water loss. According to our data, the weight loss in the beta-hex knockout mutant fruits showed no significant difference compared to the wild type during the initial three days. However, from day 6 through day 12, a significant reduction in weight loss was observed in the mutant lines compared to the wild-type controls (Fig. 2A). These quantitative findings were further corroborated by phenotypic observations; the mutant fruits exhibited superior post-harvest resilience, maintaining their firmness and structural integrity for up to 25 days, whereas the wild-type fruits showed visible shriveling and loss of turgor (Fig. 2B).

**Fig 2.**
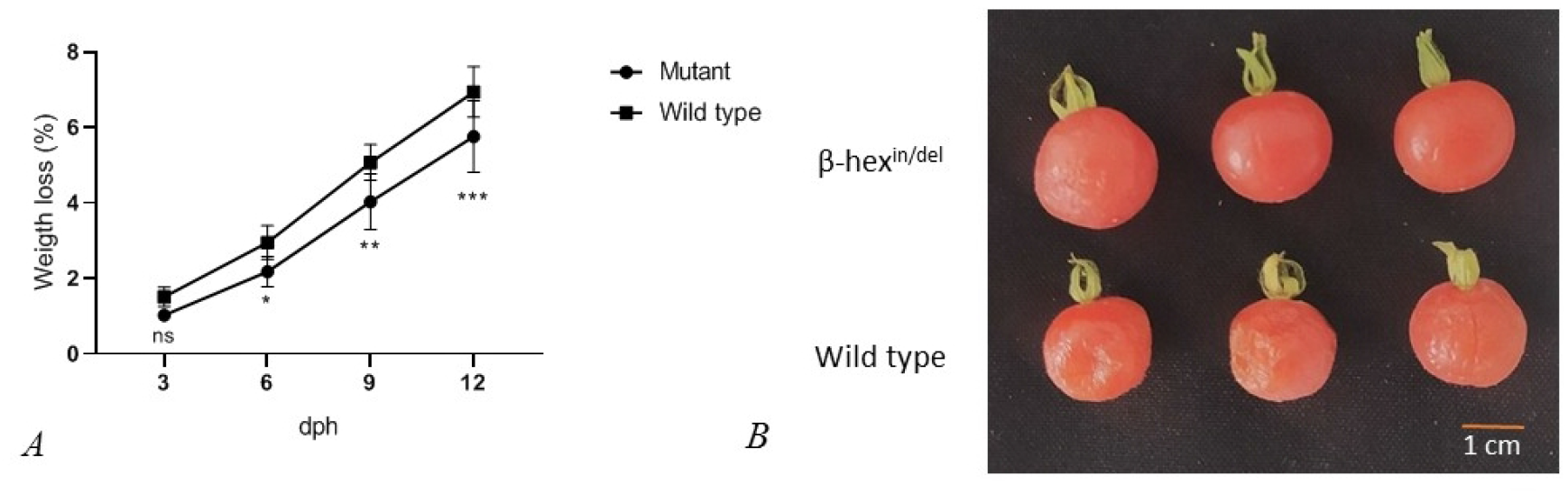
Comparative analysis of post-harvest weight loss and fruit morphology in wild-type and beta-hex mutant tomato lines. (A) Cumulative weight loss (%) of tomato fruits measured at 3-day intervals over a 12-day storage period at room temperature. Each data point represents the mean ± SD of three biological replicates. Asterisks denote statistical significance according to Student’s t-test: *p < 0.05, **p < 0.01, ***p < 0.001; ns, non-significant. (B) Representative phenotypic images of beta-hex mutant and wild-type (WT) fruits after 25 days of storage, illustrating the visible differences in fruit turgor and surface shriveling. Scale bar = 1 cm.

Fruit weight (FW), fruit shape (FS), fruit weight loss and fruit pH were measured in WT and β-hex^in/del^ plants at the mature red stage. No significant differences were detected in WT and mutant plants for FW, FS, pH characteristics (Fig. 2). However, the fruit weight loss was significantly lower in mutant plants compared to WT (Fig. 3).

**Fig 3.**
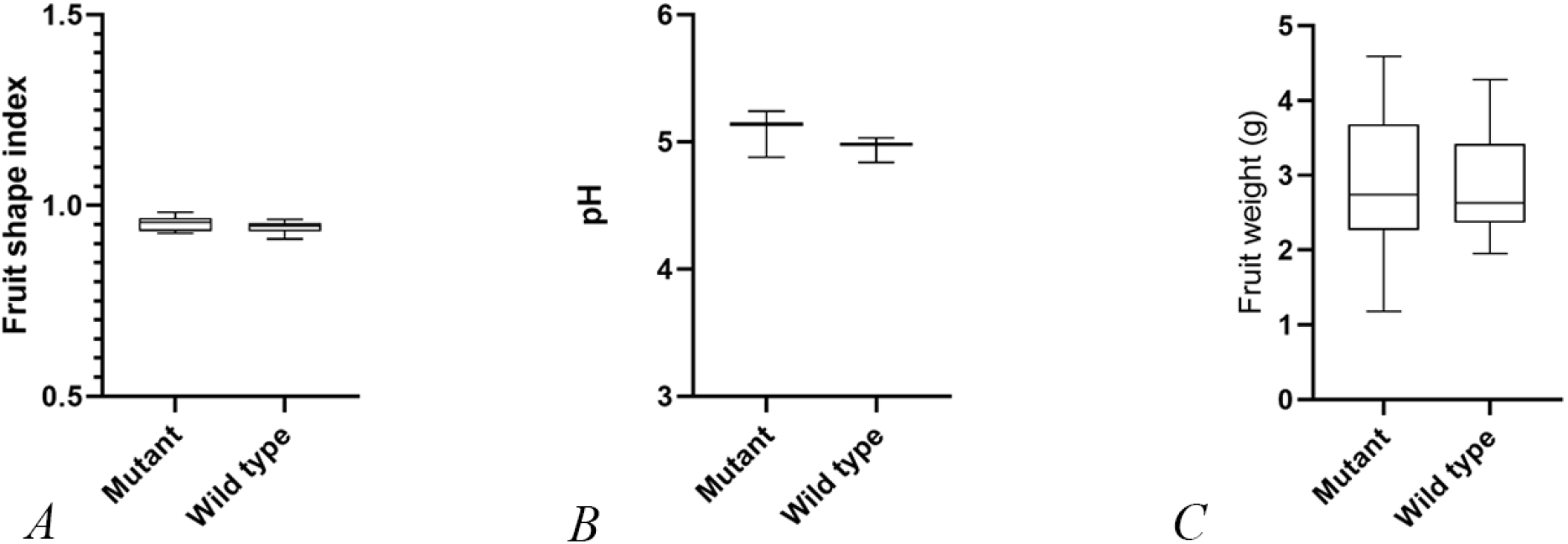
Fruit quality indicators. A fruit shape index; B pH; and C Fruit weight.

Additionally, it was found that the mutant plant’s fruits turned red much more slowly than those of the WT plant when comparing the color change of the fruits during the Breaker (Br) stage (Fig. 4).

**Fig 4.**
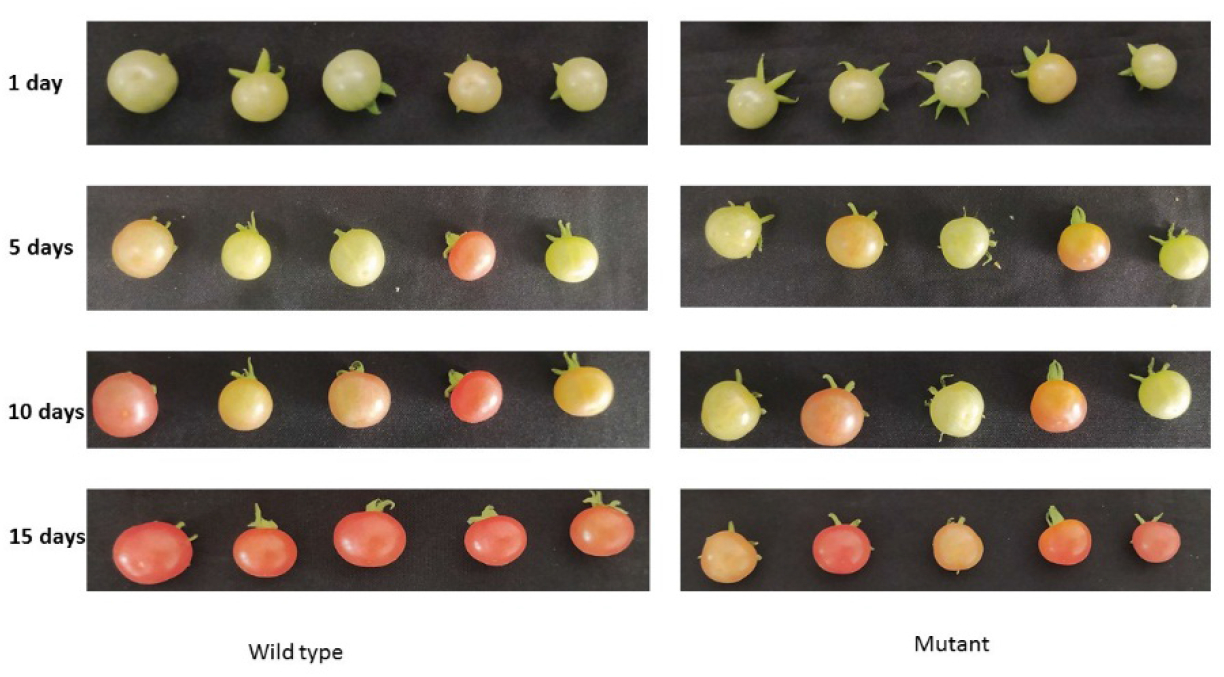
Phenotypic comparison of fruit ripening between wild-type and mutant tomato lines over 15 days. β-hex^in/del^ and Wild type fruits were harvested at breaker stage and stored at room temperature (22–24°Cin 55–60% relative humidity). The progression of fruit color changes was recorded by time-lapse photography. Time after harvest is specified by days.

## Discussion

In nature, there are NOR, Nr and CNR lines of tomato fruit with impaired ripening, and such plants can keep their fruit for a long time with good appearance [32], but it has been found that the taste of the fruit of such plants has changed. Such plants are caused by a disruption in the biosynthesis of the hormone ethylene or the ethylene receptor. Ethylene not only controls fruit ripening in the plant, but also the accumulation of nutrients in the fruit. As a result, the fruit quality of such plants is lower than that of ordinary tomatoes [32–34]. In tomato biotechnology, extending shelf life also involves a trade-off known as pleiotropic effects. For instance, editing master ripening regulators like RIN or NOR genes significantly delays ripening but results in severe defects, including inhibited lycopene accumulation and diminished flavor profiles [35].

Our study focused on the targeted knockout of the β-hex gene, which encodes a specific cell-wall-modifying enzyme rather than a global regulatory factor. In contrast to master ripening regulators, targeted modification of genes directly involved in cell wall architecture does not induce adverse pleiotropic effects on fruit quality. Specifically, the CRISPR-mediated knockout of the *SlPL* gene in tomato has been reported to maintain fruit mass, shape, and pH levels comparable to wild-type controls [36]. Similar observations were documented following the RNAi-mediated silencing of the β-hex gene [24]. Consistently, our β-hex^in/del^ mutant lines exhibited no secondary negative phenotypic changes, further supporting the precision of targeting downstream enzymes to enhance shelf life without compromising fruit characteristics (Fig. 3).

Fruit ripening is a complex mechanism involving a series of coordinated enzymatic activities. Research has demonstrated that these enzymatic processes alter the morpho-physiological characteristics of fruits by modulating specific biochemical reactions within the cell [37,38]. A critical stage in this process is the degradation of cell wall polysaccharides, such as cellulose, pectin, and glycan, by various endogenous enzymes. For instance, reducing the activity of the polygalacturonase (PG) enzyme by 40% has been shown to significantly improve fruit firmness [39]. Similarly, the suppression of β-galactosidase 4 (TBG4) activity in tomatoes using antisense technology resulted in enhanced fruit integrity compared to the control [40]. Consistently, our study on β-hex knockout lines demonstrates that modulating specific glycosyl hydrolases can significantly improve fruit firmness without disrupting essential ripening processes, such as pigment accumulation (Fig. 2B)

Plant genome editing mediated by the CRISPR/Cas9 system frequently results in predictable mutations, such as 1 bp insertions or small deletions (up to 10 bp) at the target site [41]. Similar indel mutation patterns have been extensively documented in tomato genome editing studies [42]. In our case, a 1 bp insertion in exon 1 and a 5 bp deletion in exon 2 were observed in the β-hex gene of the β-hex^in/del^ plant.

Regarding the inheritance and stability of the induced mutations, our molecular analysis showed that the T_0 primary transformants possessed heterozygous mutations at both target sites. In subsequent generations, these mutations underwent Mendelian segregation, allowing for the selection of homozygous mutant lines in the T_2 populations. Furthermore, the T-DNA construct was found to have segregated out in a significant proportion of these advanced lines (S2 Fig). The recovery of homozygous, transgene-free individuals is a crucial outcome, as it demonstrates the potential for developing genetically stable tomato varieties that lack foreign DNA, thereby addressing regulatory and consumer concerns regarding genetically modified organisms.

## Acknowledgement

We are grateful to the research team at the Center of Genomic and Bioinformatics for their assistance in interpreting the study data included in this paper. The authors would like to express their gratitude to Dr. Iris Valerie Olga Hoffie from the Plant Reproductive Biology group, Leibniz Institute of Plant Genetics and Crop Plant Research (IPK) Gatersleben, Germany, for kindly providing the CasCade vector system used in this study.

## Authors’ contributions

AAM conducted all research experiments and wrote the manuscript. MSA conceptualized the experimental design and assisted in writing the manuscript. MKM performed molecular analyses and contributed to the critical analysis of the manuscript. ANY was responsible for data analysis and the statistical part of the study. NSO, BOM, and SOK assisted in plant cultivation, growth maintenance, and phenotypic data collection. LKK assisted in the plant transformation experiments. ZHB provided assistance with statistical data processing. IEB contributed to the molecular analysis procedures. ZTB and IYA performed rigorous analysis, critically revised the manuscript, and approved the final version for publication.

